# Quantifying and engineering mucus adhesion of probiotics

**DOI:** 10.1101/731505

**Authors:** Zachary J. S. Mays, Todd C. Chappell, Nikhil U. Nair

## Abstract

Mucus in the gastrointestinal (GI) tract is the primary point-of-interaction between humans and their gut microbiota. This not only intimates that mucus ensures protection against endogenous and exogenous opportunists but provision for the human microbiota to reside and flourish. With the emergence of living therapeutics, engineered microbes can deliver and produce increasingly complex medicine, and controlling the mucoadhesive properties of different microbial chassis can dictate dose-response in a patient. Here we present a redesigned, *in vitro*, plate-based assay to measure the mucus adhesion of various probiotics. Cell-mucus interactions were isolated by immobilizing mucus to the plate surface. Binding parameters were derived for each probiotic strain by measuring cell adhesion over a wide range of cell concentrations, providing dose-dependent adhesion metrics. Surface proteins and cell components known to influence mucoadhesion were then heterologously expressed or altered in *Lactococcus lactis* MG1363 and *E. coli* Nissle 1917 to control mucus-binding capacity, avidity, and cooperativity.

---

As a new era of precision medicine and personalized therapeutics becomes increasingly main-stream, there too is heightened interest in our ability to leverage the human gut microbiota, a complex and rich community of approximately 1000 species of microbes living in the gastrointestinal (GI) tract. Although causality remains uncertain, recent evidence suggests a strong relationship between many disease states and a perturbed, dysbiotic, gut microbiota^1, 2^. One strategy to restore microbial balance, and the basis of 282 clinical trials in the last decade^3^, is to introduce probiotics – microbes shown to confer health benefits. Probiotics have been shown to inhibit or exclude enteric pathogens^4, 5^, modulate or improve mucosal barrier function^6^, reduce inflammatory responses^7^, ameliorate metabolic or toxin imbalances^8^, maintain intestinal pH, and contribute to neurohumoral signaling^9^. Furthermore, engineered probiotics as therapeutic delivery vehicles^10–12^ are being explored for pathogen eradication^13^, vaccines^14^, metabolic disorders, enzyme-replacement therapies^15, 16^, cancer treatments^17, 18^, and diagnostics^19, 20^.

Somewhat counterintuitively, the stability of a healthy gut microbiota likely arises from its diversity and competitiveness rather than established cooperative metabolic networks^21^. Thus, an argument can be made that when developing probiotic therapeutics, one design focus should be to enhance factors conferring competitive opportunity (of course, while safeguarding against overgrowth). This dichotomy is further exemplified by the fact that lactic acid bacteria (LAB), which encompass the majority of common probiotics studied for disease amelioration, make up less than 1% of the human microbiota^22^. Furthermore, orally-administered probiotics are rarely detected past a week after ingestion, and persistence is highly strain- and patient-specific^23^, making dose-response relationships unreliable^24^. From an engineering perspective, directing a microbe toward a synthetic application inherently places a burden that conciliates competitive advantage compared to its natural counterpart in the body; essentially, current strategies cannot persist, or even survive, long enough to be effective.

The most prominent point-of-interaction between microbes and humans is at mucus membranes, revealing mucus adhesion as a prime target for controlling probiotic occupancy. Indeed, the cell-surface structures of commensal bacteria are hypothesized to have coevolved in the presence of mucus glycans, creating preferential mucoadhesive mechanisms^25^. Additionally, the extracellular polysaccharides (EPS) secreted for LAB biofilm formation can influence colonization through mucus associations^26, 27^, while pathogenic biofilm formation is impeded by LAB extracellular polysaccharides and mucus itself^28^. Mucin has also been observed to increase surface-protease expression in LAB, suggesting the utilization of mucus as a nutrient source^29^. Despite these interactions and the mucus matrix of the GI tract providing a topography for microbes to find spatial niches, orally-administered natural probiotics and engineered microbial therapeutics do not currently provide any means of controlling biogeographical targeting, residence time, or an ability to colonize the gut. The ability to accurately measure and engineer mucus adhesion could address these limitations and provide a new avenue to increase efficacy of natural and engineered probiotic therapies.

Methodologies to quantify bacterial adhesion to mucus have been adapted from traditional cell adhesion assays and inherit similar advantages and disadvantages^30^. The general procedure behind these adhesion assays is to quantify the fraction of residual bacteria bound to mucus-functionalized surfaces after washes^31^. Quantifying residual bacteria can range from a simple cell count^32^ to surface plasmon resonance^33^ or atomic force microscopy^34^. The most common method to quantify mucus adhesion is to use a fluorescent indicator as a correlate for cell concentration. Surfaces are often modified by incubating with mucus or by culturing gut epithelial cell/organ tissue. However, recent studies showed that bacteria adhere to polystyrene, the support of mucus functionalization, as well or better than mucin- or cell culture-coated microplate wells^35^. Additionally, there are several examples of bacterial strains showing different adhesion characteristics depending on the mucus source^36–38^, and different mucus compositional profiles can be observed in co-culture models simply by differentiating the HT29 cell line with either methotrexate (MTX) or 5-fluorouracil (FU)^39^. The question arises then, whether these adhesion data are the result of the mucus composition affecting cell-cell associations, cell-mucus associations, or mucus-plastic associations. Measuring more holistic binding parameters when quantifying mucus adhesion would allow for better comparability across conditions.

In order to address these disconnects, we report an improved *in vitro* method for quantifying bacterial adhesion to mucus. Rather than relying on natural nonspecific interactions to coat a microplate with mucus, we covalently functionalized surfaces with mucus, providing robust and full coverage. Our assay is able to isolate the cell adhesion events associated directly with mucus. We then measure adhesion over a range of cell concentrations to develop strain-specific mucus-binding curves. By more clearly describing mucus adhesion using strain-specific quantifiers interpolated from a mucus binding curve, we can better compare various binding characteristics related to binding capacity, binding avidity, and cooperativity for common probiotic strains as well as common probiotic engineering strains. We also demonstrate how the expression of recombinant characterized and putative mucus-binding proteins, or deletion thereof, can be used to alter mucus-binding characteristics of probiotics and how these changes are reflected in the binding curves. Such altered mucus-binding has never been engineered previously. Overall, our methods and results provide a novel and robust approach to quantifying bacterial adhesion to mucus and advances design of microbial therapeutics with tunable pharmacological release factors.

## RESULTS AND DISCUSSION

### Silanization and EDC-coupling covalently binds mucus to the plate surface

A comparison of traditional *in vitro* models used to study bacterial adhesion to the intestinal epithelium revealed that differences between bacterial strains binding to mucus are significantly confounded by their ability to strongly stick to the abiotic surface of the microplate^35^. To remedy this, we sought to covalently link mucus to the plate, providing full mucus coverage and removing cell-plate interactions. To perform this, the microplate surface was silanized with (3-aminopropyl)triethoxysilane (APTES) to provide a surface of available primary amine groups. The most abundant gel-forming mucin protein in the mammalian small intestine is Muc2, which is contains sialylated and sulfated terminal glycans that contain freely accessible carboxyl groups^40, 41^. Using the primary amine on the plate and available carboxyl groups from the terminal sialic acids of Muc2, mucus was then crosslinked to the silanized microplate using a common *N*-ethyl-*N*’-(3-dimethyl-aminopropyl) carbodiimide and *N*-hydroxysuccinimide (EDC/NHS) reaction mixture (Figure 1). Porcine intestinal mucus (PIM) was used as an analog for human mucus because of the compositional and rheological similarities^42, 43^. Though mammalian mucus glycobiology is diverse and species specific, core 3-derived *O*-glycans are relatively consistent (Figure 1Figure 1^40^ and are the major glycan structure of Muc2 in the mammalian intestine^44^.

**Figure 1.**
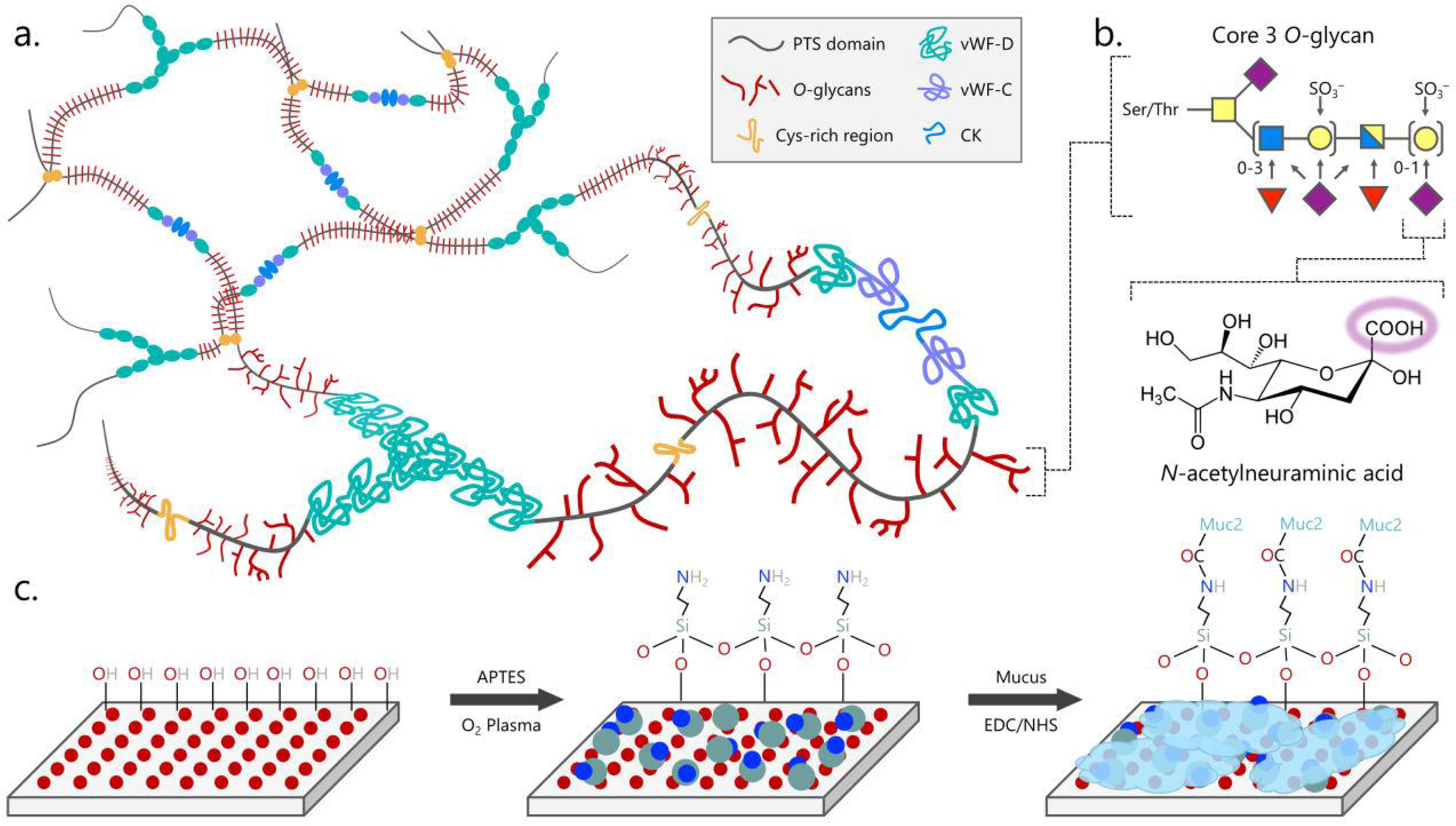
Schematic of mucus immobilization strategy. The structure of mucus presents an opportunity for covalent immobilization chemistry. Gel-forming mucins such as Muc2 form (a.) macromolecular sheets of trimers of dimers that crosslink or interact to form a mesh between von Willebrand Factor (vWF) domains and cysteine knots (CK). Each mucin monomer has a highly glycosylated proline-threonine/serine (PTS) domain with the (b.) highly sialylated core 3 O-glycan dominating in the intestines. Sialic acid, or N-acetylneuraminic acid (purple diamond), provides a freely available carboxylic acid that can be (c.) coupled to a primary amine presented on a glass or plastic surface using EDC/NHS crosslinking chemistry. Other common glycans include *N*-acetylglucosamine (blue square), *N*-acetylgalactosamine (yellow square), galactose (yellow circle), and fucose (red triangle). *Final optimized conditions for full mucus coverage require 1% PIM, 10x higher than other conditions, and ELISA luminescence values increased proportionally.

Previous reports indicate that it is possible to immobilize mucus using EDC-coupling, wherein mucus provides primary amine groups for the reaction^33^. Because our proposed reaction scheme requires mucin to donate carboxyl groups, we wanted to confirm that EDC-coupling was indeed possible with the plate surface and that the reaction was not hindered by a self-reaction involving amines within the mucins themselves. To accomplish this, we performed two EDC reactions in tandem. First, between fluorescent sulfo-cyanine5 amine (cya5) and PIM in which mucus provides carboxyl groups and a subsequent filtration removes any unbound fluorophore. Second, between PIM and the microplate in which vigorous wash steps can separate non-specifically bound mucus from that which is covalently attached to the plate. Fluorescence was observed before and after washing, and the presence of mucus was confirmed by a follow-up Muc2-specific enzyme-linked immunosorbent assay (ELISA) (Figure 2a). As shown in Figure 2b, when both EDC-coupling reactions are performed, not only is fluorescence observed initially, suggesting covalent linking to the mucus, but it remains after multiple washes, suggesting the mucus is stably linked to the microplate surface. Absence of the EDC reagent at either stage results in no fluorescence or plate binding, as expected. Similar results were observed from the subsequent Muc2-specific ELISA (Figure 2c). These results confirm the directionality of the EDC-coupling reaction and that mucus was covalently bound to the microtiter plate. Covalently bound mucus provided further opportunities for optimizing the cell adhesion assay, including automated wash steps and added shear stress during the incubation using orbital shaking.

**Figure 2.**
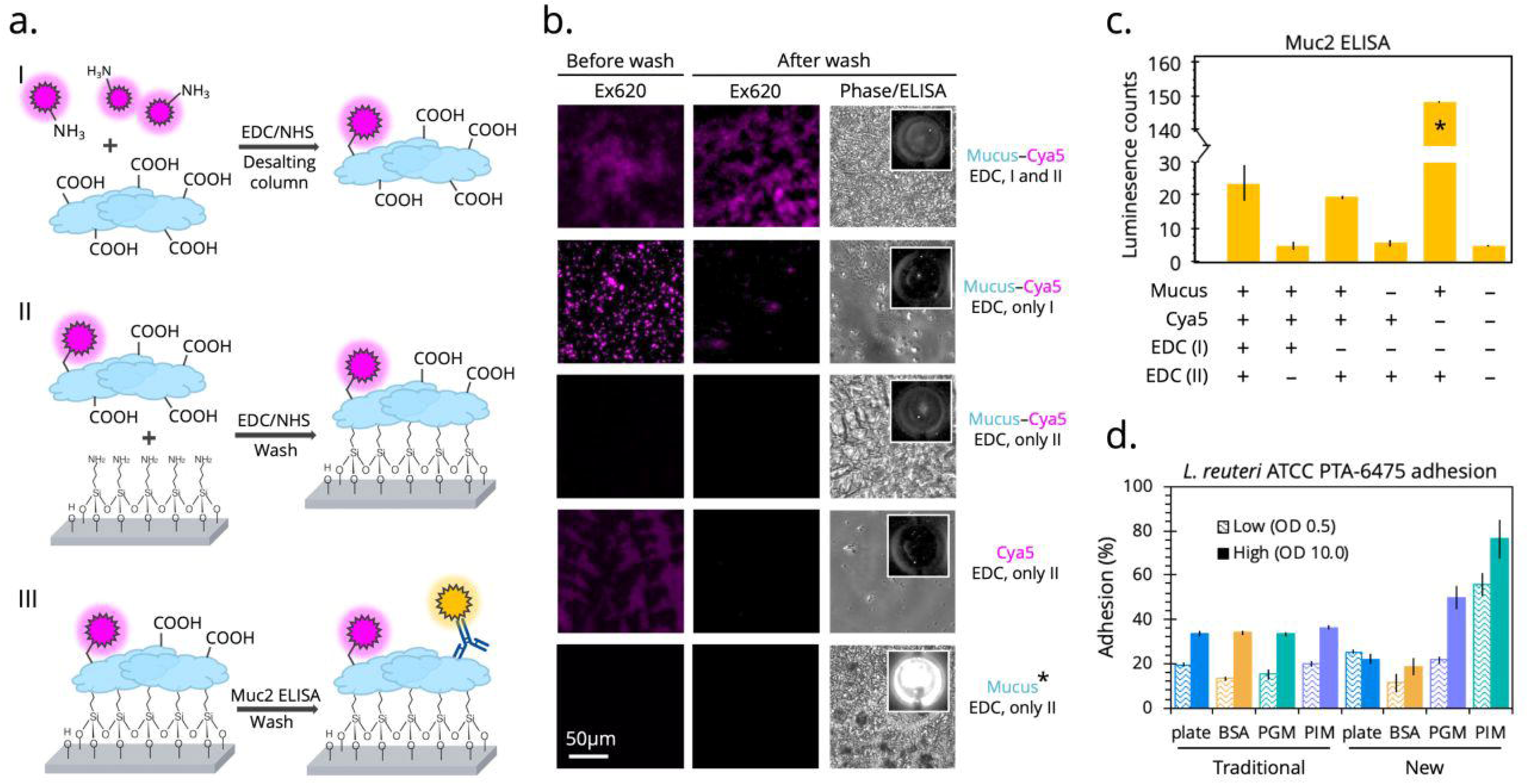
Validating the modified cell mucoadhesion assay and comparing to the traditional assay. (a.) Two EDC/NHS reactions were performed in tandem followed by a Muc2-specific ELISA to determine the success of the mucus immobilization. (b.) Fluorescence microscopy was then implemented to measure of the robustness and directionality of the coupling between mucus and the sulfo-cyanine5 fluorophore and between mucus and the plate surface. Each condition was imaged before and after vigorous washing and after the ELISA. (c.) Luminescence counts were quantified to determine mucus coverage, with the final optimized condition (1% PIM) measuring 10-fold higher than other conditions, as expected. (d.) With these final conditions, the traditional adhesion assay was compared to the newly op-timized assay using the strong binder *L. reuteri* ATCC PTA-6475 as a metric for success on different protein surfaces. Percent adhesion was calculated from fluorescent output of cFDA-stained cells.

### Modified microtiter plate assay reveals dependence of cell adhesion on mucus type

Bacterial strains isolated from mammalian intestines have been reported to preferentially bind mucus over other general proteins such as bovine serum albumin (BSA)^38^. Unfortunately, when using the traditional cell adhesion assay to test the mucoadhesive ability of *Lactobacillus reuteri* ATCC PTA-6475 at OD_600_ 0.5, we were not able replicate preferential binding to mucus over BSA (Figure 2d). Only after implementing our newly modified covalent immobilization, did we observe significant differences between binding to polystyrene plate, BSA, porcine gastric mucus (PGM), and PIM by *L. reuteri* ATCC PTA-6475. Specifically, there was a larger percent of adhered *L. reuteri* ATCC PTA-6475 cells to PIM than the plate, BSA, and PGM (Figure 2d), potentially recapitulating this strain’s co-evolution with, and preferential binding to, intestinal niches.

It has been previously reported that mucus gel reconstituted from commercially available PGM does not accurately replicate the pH-dependent rheological characteristics (viscosity and elasticity) of native mucus^45^ and has an altered structure, likely from industrial processing^46^. Polymer rigidity, influenced by the negative charges of sulfates and terminal sialic acids, is an important determinant of the viscosity and elasticity of mucus^47^, which may explain the diminished cell adhesion capacity observed with the processed PGM (Figure 2d). Maintaining the glycan chemical identity of mucus is more important than macromolecular structural integrity when interrogating mucus adhesion. This is highlighted by the fact that a common mucoadhesion assay is to separate, purify, and freeze mucins to be later used in dot-blot assays in which no mucus gel is required^48^. As shown in Figure 2b, Muc2 antibody recognizes the covalently attached mucus indicating that important epitopes are preserved through our method processing.

We obtained similar preferential binding to covalently immobilized PIM at a high cell density (OD_600_ 10.0) of *L. reuteri* ATCC PTA-6475 while the traditional assay demonstrated equivalent and non-specific maximum binding capacity of 35% under all conditions (Figure 2d). This was likely observed because the more vigorous washing required at higher cell loadings removes non-covalently attached proteins from the plastic surface. Additionally, while *L. reuteri* ATCC PTA-6475 did not appear to bind any better to PGM than BSA or the plate at the low cell density, the binding capacity significantly increased at the higher cell density, becoming higher than the 35% maximum threshold of the uncoat-ed plate (Figure 2d). It can then be inferred that covalently immobilized mucus enables detection of specific cell-mucus interactions rather than non-specific cell-plastic binding events that dominate on non-covalently functionalized plates. A final important aspect of the new assay is a higher throughput and automated wash procedure, adding consistency between technician operability without sacrificing replicate variability.

### Different probiotics display vastly different mucoadhesive characteristics

Cell mucoadhesion is typically reported as a percent or fractional residual bound cells at a single cell concentration, often at OD_600_ 0.5^37, 38, 48–51^. To assess binding characteristics in greater resolution, we measured adhesion of several probiotic strains and species across a wide range of cell concentrations (OD_600_ 0.01 to 100). Adherence was calculated by staining the cells with the fluorescent indicator carboxyfluorescein diacetate (cFDA) and detecting the fluorescence at each concentration before and after washes. The dye only fluoresces when taken up by the cell and deacetylated by native esterases. In order to avoid non-specific surface binding that may affect cell mucoadhesion, we stained the cells at a much lower concentration than typically used^52^ (10 μM versus 100 μM). There is no observed cytotoxicity at this concentration^53^, a sign that membrane integrity is maintained.

While there are some previous reports measuring dose-dependent adhesion of probiotics, these were either performed using a Caco2 monolayer^54^, therein not directly measuring mucus adhesion, or performed at very low cell concentrations (OD_600_ < 0.1), likely due to the limited dynamic range of the traditional assay^55^. With our more robust immobilized mucus plate, we aimed to develop a more quantitative and descriptive mucoadhesive measurement by expanding the dose range, introducing shear stress in the form of orbital plate shaking, and fitting a sigmoidal curve to the adhesion data to make direct comparisons without the need to normalize between strains. With many cell-mucus interactions determined to be lectin-like recognition of mucus glycans by surface proteins^56–59^, a sigmoidal Hill-type shape^60^ was chosen with the hypothesis that cell-mucus binding mirrors the energetics of ligand-receptor binding.

Using this method, distinct strain-specific differences between the mucoadhesive abilities of common probiotics become apparent (Figure 3). This was particularly interesting for *L. reuteri* human isolates. *L. reuteri* ATCC PTA-6475 and *L. reuteri* ATCC PTA-5289, which are nearly identical at the genome level^61^ clustered in the MLSA (multilocus sequence analysis) lineage II phylogenic group^62^. Notably, these strains also contain variations of cell and mucus-binding protein A (CmbA)^49^ and have been previously reported to have strong adhesion to mucus and intestinal epithelial cells (IEC)^63^. In contrast, *L. reuteri* DSM 17938 is from a different phylogenic branch (lineage VI) and has a much lower reported adherence to IECs^48^, which concurs with our observation that it attains a low maximum adherence capacity (A_max_ = 4.3×10^7^ cells/cm^2^).

**Figure 3.**
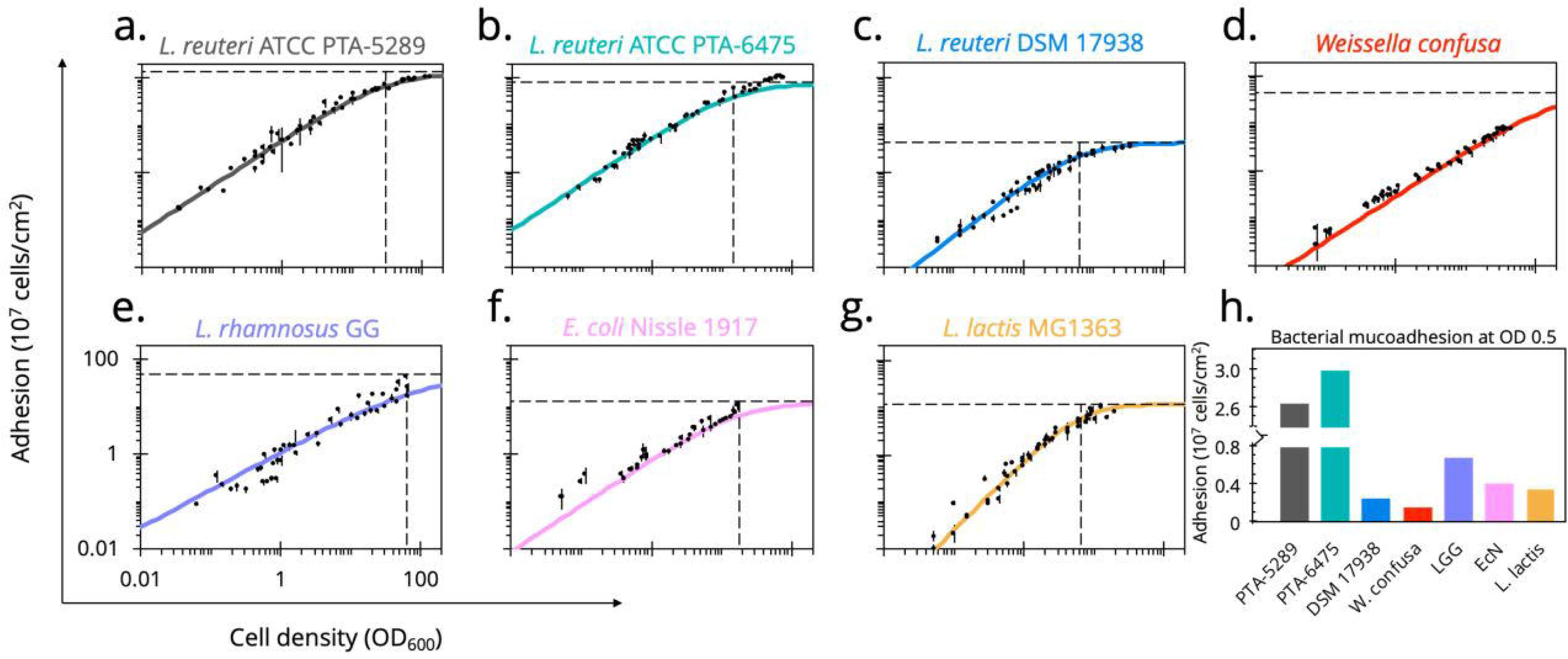
Dose-dependent mucus-binding curves for common probiotics. (a.-g.) Using our optimized plate assay, the number of cells adhered to immobilized mucus was measured over a large range of concentrations for common probiotics. Data points are reported as the mean values of quadruplicate wells across three independent experiments. Hill-type sigmoidal curves were fit, with curve parameters a metric for strain-specific mucoadhesive characteristics (Table 1). (h.) With the traditional assay typically performed at OD600 0.5, comparative analysis across strains was visualized at this density.

*Escherichia coli* Nissle 1917 (EcN) is an extensively studied probiotic sold over-the-counter as Mutaflor® to treat chronic inflammatory and infectious intestinal diseases and is used significantly in the development of microbial therapeutics^16, 64–69^. Though there are conflicting reports whether EcN can persist in human intestines^68, 70^, it has been shown to colonize the porcine intestinal tract^71^. Therefore, our results indicate a surprisingly poor ability to adhere to PIM, as determined by this assay (A_max_ 12.9×10^7^ cells/cm^2^). Like *L. reuteri* DSM 17938, its binding plateaus to its maximum relatively easily, reaching half maximum adhesion at OD_A50_ 17.7 (Table 1). It has been suggested that biofilm-like aggregation is necessary for EcN colonization^72^ and that mucoadhesion is influential in initial binding before other competitive mechanisms take over. Additionally, compared to pathogenic strains, commensal *E. coli* have more flexibility in their ability to perform metabolic shifts depending on the mono- and disaccharide carbon sources available from mucin glycans^73^. This suggests commensal *E. coli*, such as EcN, may utilize mucus by finding a nutritional niche rather than adhering as a persistence mechanism.

**Table 1.** Summary of mucus-binding curve parameters for common probiotics

| Strain | <sup>a</sup>A<sub>max</sub><br>(×10<sup>7</sup> cells/cm<sup>2</sup>) | <sup>b</sup>OD<sub>A50</sub> | n |
| --- | --- | --- | --- |
| *L. reuteri* ATCC PTA-6475 | 77.4 ± 3.1 | 14.3 ± 1.5 | 0.98 ± 0.05 |
| *L. reuteri* ATCC PTA-5289 | 133.5 ± 7.5 | 30.0 ± 4.1 | 0.97 ± 0.05 |
| *L. reuteri* DSM 17938 | 4.3 ± 0.3 | 6.2 ± 1.1 | 1.11 ± 0.11 |
| *Weissella confusa* | 44.7 ± 0.0 | <sup>a</sup>203.7 ± 0.1 | 0.95 ± 0.001 |
| *L. rhamnosus* GG | 44.5 ± 0.3 | <sup>a</sup>99.0 ± 0.8 | 0.80 ± 0.02 |
| *E. coli* Nissle 1917 | 12.9 ± 0.2 | 17.7 ± 0.2 | 0.98 ± 0.04 |
| *L. lactis* MG1363 | 6.8 ± 0.4 | 3.7 ± 0.4 | 1.52 ± 0.14 |

Parameters determined from sigmoidal curve fitting of experimental adhesion data, shown with 95% confidence intervals
<sup>a</sup>Extrapolated value from curve fitting
<sup>b</sup>Standard curves to correlate OD<sub>600</sub> to CFU/mL are reported for each strain in Table S1.

Another commonly used clinical probiotic is *Lactobacillus rhamnosus* GG (ATCC 53103), which has been reported to have strong mucoadhesion primarily through SpaCBA pilus-mediated interactions^74^. Notably, both EcN and *L. rhamnosus* GG show moderate mucus-binding ability and use moonlighting proteins with the secondary function of mucus adhesion but have quite different binding characteristics. While the probiotic nature of *Weissella confusa* is in question^75^, it was observed to have a similar binding capacity (A_max_ 45 ×10^7^ cells/cm^2^) to *L. rhamnosus* GG, which corroborates a report of these strains binding strongly to HT29-MTX, a mucus-producing differentiated epithelial cell line^76^.

Finally, *Lactococcus lactis* is undoubtedly the most well-studied LAB and the basis of centuries of food engineering. Importantly, *L. lactis* MG1363 showed poor binding ability to PIM compared to other probiotic strains (Figure 3g). Despite similar curve parameters to *L. reuteri* DSM 17938, *L. lactis* MG1363 has a higher sigmoidal coefficient, creating a tighter window of concentrations in which mucus binding is influential compared to other strains.

### Expression or removal of surface proteins can be used to alter adhesion characteristics

There are several surface proteins characterized on LAB that either moonlight or directly mediate mucoadhesion^49^. A database search revealed a number of known and putative mucus-associated proteins found on the *L. reuteri* strains being evaluated. Constitutively expressing three of these proteins in *L. lactis* MG1363 highly, moderately, and minimally altered the stain mucoadhesion characteristics.

Cell and mucus-binding protein (CmbA) has been reported to significantly impact the ability of *L. reuteri* ATCC PTA-6475 to bind to Caco-2 cells and PIM^25^. The protein also contains a 288 base pair tandem repeat region, a characteristic found in surface proteins associated with mucus binding in various LAB^77^. When expressing CmbA on the surface of *L. lactis* MG1363, the mucus-binding ability of the strain drastically improved, creating a binding curve with similar parameters to that of *L. reuteri* ATCC PTA-6475 (Figure 4a and Table 2).

**Figure 4.**
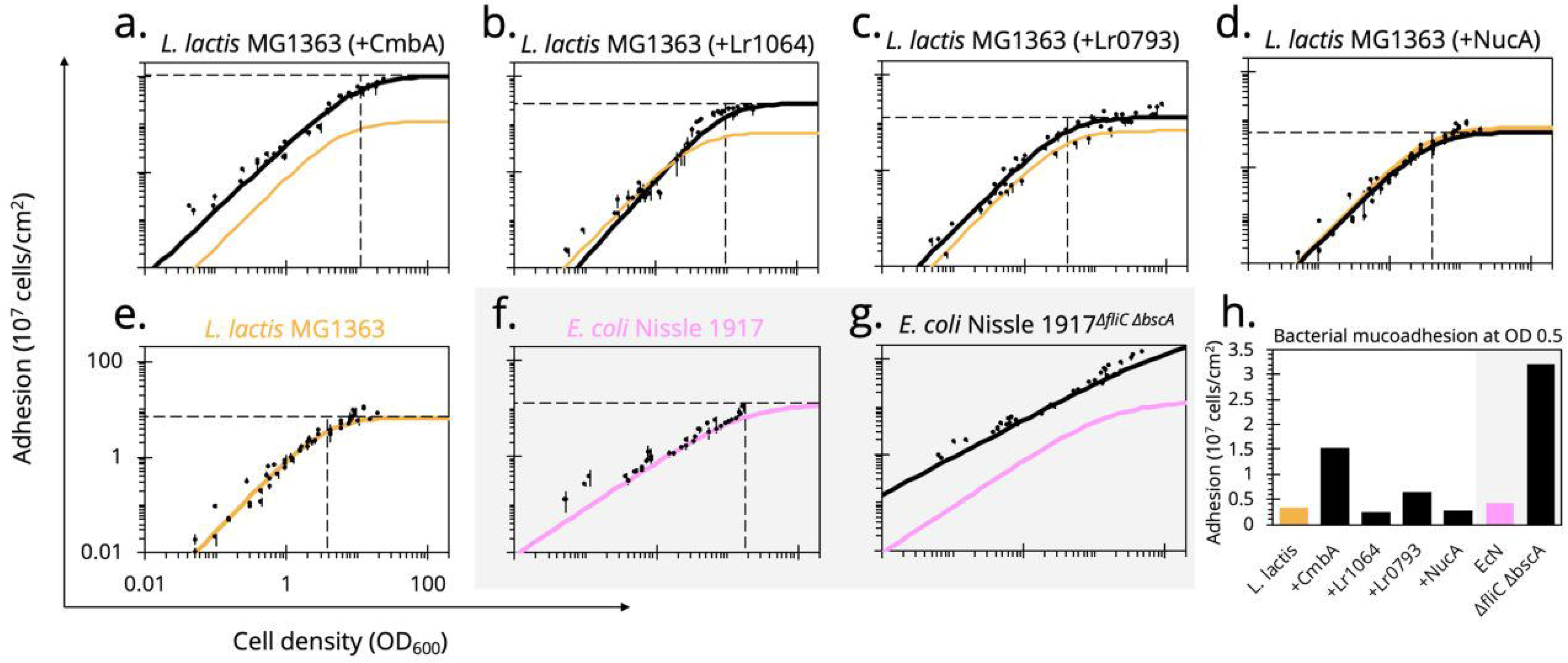
Altered dose-dependent mucoadhesion by heterologous protein expression. Known and putative mucus-associating surface proteins from *L. reuteri* were heterologously expressed in *L. lactis* MG1363. The (e.) wild-type *L. lactis* MG1363 binding curve (orange) was (a.-c.) compared to the engineered counterpart (black) as well as the *L. reuteri* strain from which the protein was sourced (green or blue). (d.) As a display control, an inactivated NucA^E41Q^ did not alter mucus binding compared to the wild-type strain. Alternatively, the flagellum of EcN, considered a known mucus-binding factor, and cellulose synthase, a contributor to biofilm formation, were (g.) knocked out but surprisingly improved mucoadhesion compared to the (f.) wild-type (pink). (h.) With the traditional assay typically performed at OD_600_ 0.5, comparative analysis across strains was visualized at this density.

**Table 2.** Summary of mucus-binding curve parameters when altering mucoadhesive protein expression

| Strain | <sup>a</sup>A<sub>max</sub><br>(× 10<sup>7</sup> cells/cm<sup>2</sup>) | OD<sub>A50</sub> | n |
| --- | --- | --- | --- |
| *L. lactis* MG1363 | 6.8 ± 0.4 | 3.7 ± 0.4 | 1.52 ± 0.14 |
| + NucA<sup>E41Q</sup> | 5.5 ± 0.4 | 4.0 ± 0.5 | 1.5 ± 0.13 |
| + CmbA | 104.3 ± 4.6 | 11.5 ± 0.8 | 1.37 ± 0.12 |
| + Lr1064 | 27.8 ± 1.7 | 9.4 ± 0.9 | 1.64 ± 0.12 |
| + Lr0793 | 13.0 ± 0.5 | 4.1 ± 0.5 | 1.48 ± 0.22 |
| *E. coli* Nissle 1917 | 12.9 ± 0.2 | 17.7 ± 0.2 | 0.98 ± 0.04 |
| ΔfliC,ΔbscA | 368.3 ± 5.5 | <sup>a</sup>230.5 ± 3.6 | 0.78 ± 0.02 |

Parameters determined from sigmoidal curve fitting of experimental adhesion data, shown with 95% confidence intervals
<sup>a</sup>Extrapolated value from curve fitting

Despite the observed poor mucus-binding ability of *L. reuteri* DSM 17938, the cell surface of *L. reuteri* SD2112, the genetically equivalent parental strain, has been characterized with multiple identified putative mucus binding proteins, including Lr0793 and Lr1064^78, 79^. The protein Lr0793 is an ATP-binding cassette (ABC)-transporter component of an amino acid transporter, homologous to collagen-binding protein (CnBP) and mucus adhesion promoting protein A (MapA) with 97% sequence identity. These homologs have been described as binding to collagen type I^80^, gastric mucus^81^, and Caco-2 cells^77^. However, expressing Lr0793 on the surface of *L. lactis* MG1363 only slightly improved maximum adherence capacity to PIM compared to wild-type while OD_A50_ remained unchanged (Figure 4c and Table 2). Similarly, Lr1064 is an ABC-transporter component of a metal ion transporter and member of the LraI family^82^. These proteins have been reported to be adhesins of oral streptococci^77^. When expressed on *L. lactis* MG1363, Lr1064 increased the maximum adherence capacity (27.8×10^7^ cells/cm^2^) while keeping the OD_A50_ similar to wild-type, thus moderately improving mucoadhesion (Figure 4b and Table 2). Additionally, when testing the binding ability of *L. lactis* MG1363 expressing Lr1064 on BSA rather than PIM, there was no observed improved adhesion, reinforcing the observed differences are in fact mucus-specific cell adhesion caused from surface protein expression (Figure S1a). This further emphasizes that our processing does not alter mucin glycans necessary for cell adhesion. These studies demonstrate the binding of recombinant *L. lactis* recapitulates that of the strain from which the mucus binding proteins were sourced.

Expression of each mucus-associated protein was confirmed by Western Blot (Figure S2), and surface display was confirmed by flow cytometry (Figure S3) using anti-His-tag antibodies. The pSIP expression vectors have previously shown successful secretion of the reporter nuclease A (NucA) in LAB^83^. We displayed an inactive mutant (NucA^E41Q^) on *L. lactis,* as a display control, and confirmed that this strain demonstrated no improvement in mucus binding (Figure 4d), again reiterating that the *L. reuteri* proteins enhance binding in a mucus-specific manner.

In consideration of altering mucus adhesion to increase probiotic residence time in the gut, the engineered *L. lactis* MG1363 strains were more resilient against washout on the plate than the wild-type counterpart. By measuring fluorescence over successive washes, strong mucus binders, such as *L. reuteri* ATCC PTA-6475, not only adhere at a higher initial density, but they remain bound over multiple applications of shear stress (Figure 5). This resilience is also recapitulated to different degrees in the engineered *L. lactis* MG1363 strains, indicated by a less negative slope over the washing steps (i.e. fewer cells removed compared to wild-type *L. lactis* MG1363). The improvement is more prominent at a low cell density of OD_600_ 0.5 compared to OD_600_ 5.0 (Figure 5), which may be a result of saturating the mucus binding sites available on the plate at higher cell densities. Lactic acid bacteria are some of the few species that can survive in the harsh conditions of the stomach and small intestine, resulting in low microbial biomass (10^2^-10^5^ cells/mL) in this section of the GI tract^84^. Our results suggest residence time of delivered probiotics or microbial biotherapeutics can be increased in the small intestines without necessarily requiring competitive binding or displacement. We have performed a preliminary competition assay, testing the ability of a strong mucus binder to displace a poor mucus binder, and again only observe competition only at very high cell densities (Figure S4).

**Figure 5.**
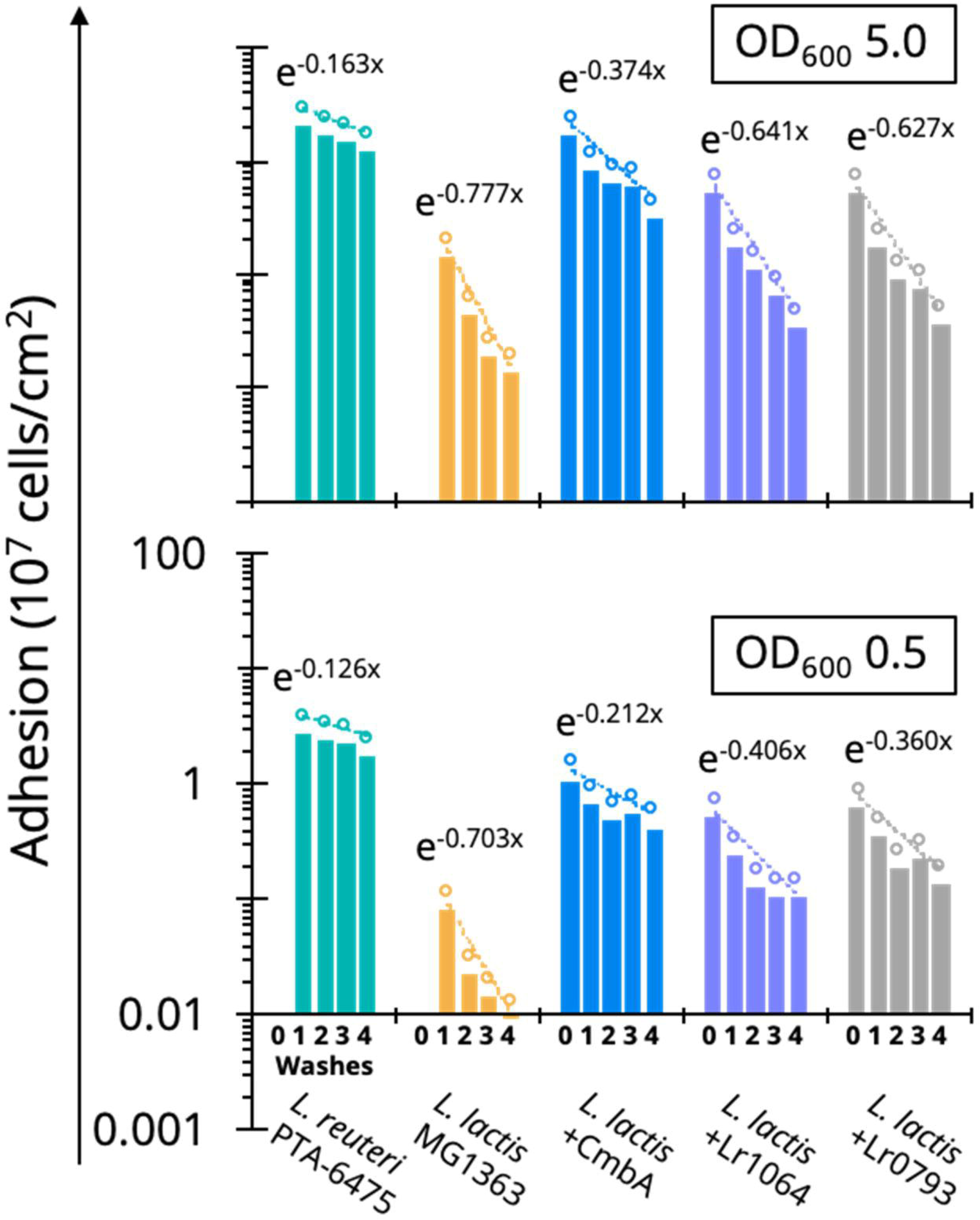
Mucoadhesive resilience against washout for engineered LAB. By observing the remaining fluorescent cells adhered to the plate over successive washes, a washout factor can be calculated from the trend (the exponential slope). The less negative the slope, the fewer cells are washed away each round, suggesting that not only can strong mucus binders (*L. reuteri* ATCC PTA-6475) adhere at higher initial density, but they are more resilient in the presence of shear stress. Resilience can be engineered by expressing mucus-adhesion factors on the surface of a poor mucus binder (*L. lactis* MG1363). Resilience against washout is also more apparent at a low cell density (OD_600_ 0.5) when mucus binding sites are not saturated on the microtiter plate.

While mucus adhesion influences residence time and biogeography, the primary selective pressure in the gut is likely centered around metabolism. For example, in a mouse microbiota with low diversity (associated with dysbiosis), EcN was observed to shift from gluconate to *N*-acetylglucosamine metabolism as a stress response^85^. Additionally, commensal *E. coli* are more flexible and sequential in their carbon utilization while pathogenic *E. coli* consume multiple carbon sources simultaneously in a slash and burn approach, highlighting the different strategies between scavenger and infection systems in the gut^73^.

The major mucus adhesin on EcN is considered to be the flagellum, formed with the structural subunit FliC^86^. Additionally, EcN expresses a cellulose synthase (BscA) necessary for biofilm formation and adhesion to epithelial cells^87^. We sought to assess the role of FliC and BcsA in EcN mucus binding. Somewhat unexpectedly, and counter to previous reports, deleting both genes (EcN^*ΔfliC,ΔbscA*^) increase the mucoadhesive ability of EcN (Figure 4g), shown by a curve shift up and to the left. This relationship is reversed, however, when the adhesion of each strain was subsequently tested on polystyrene alone (Figure S1b). Recently, the *E. coli* flagellum was shown to adhere more tightly to hydrophobic abiotic surfaces than hydrophilic substrates and may in fact impede other surface adhesins as a tradeoff for mobility^88^. This behavior is consistent with our results in Figure 4g, suggesting the adhesion of EcN to the hydrophilic mucus surface was improved by removing the flagellum, and it is possible that previous measurements of EcN mucoadhesion were either confounded by the cells binding other available abiotic surfaces like plastic. These results emphasize the importance of covalently immobilizing mucus and measuring global binding parameters across a wide range of cell concentrations. Beyond the initial observation that more EcN^*ΔfliC,ΔbscA*^ cells remain bound than EcN, the binding parameters show a stark difference in curve shape between the wild-type and knockout strains, perhaps suggesting different mechanisms of adhesion (Table 2). This is further emphasized when observing binding on polystyrene alone and with the observation that the removal of the flagellum improves binding to PIM and BSA indiscriminately (Figure S1). Despite a 46% increase in adhesion at OD_600_0.5, EcN^*ΔfliC,ΔbscA*^ retains a similar curve shape to EcN because nonspecific interactions dominate the adhesion of both cell types to polystyrene (Figure S1c). Curve shapes are altered when protein-protein interactions (mucus-specific or otherwise) at surfaces dominate (Figure 4h).

## CONCLUSIONS

Our improved mucus adhesion assay isolates mucus-specific interactions by immobilizing a confluent layer of mucus to a plate surface. This removes previously observed variability caused through non-specific adhesion mechanisms to the polystyrene while retaining the benefits of a minimalistic adhesion assay. The assay is further improved by implementing automated wash steps for consistency, and, measuring adhesion over a wide range of cell concentrations facilitates the interpolation of a mu-cus-binding curve. Curve parameters offer a more global metric for strain-specific mucoadhesion without the need to normalize to a reference strain. We observe clear dose-dependent differences in the binding ability of a number of probiotic strains with the curve parameters extending these relationships beyond the traditional measurement of percent adhesion at one cell density. Importantly, known and putative mucus-associated factors were expressed on the surface of a poor mucus binder, *L. lactis* MG1363, and mucoadhesion was significantly altered, reinstating the mucus-specific interactions of these cell-surface elements. Furthermore, a new understanding of EcN mucoadhesion was observed with the polar flagellum, originally considered a major mucoadhesin, actually impeding mucus adhesion and more general hydrophilic surface adhesion. This emphasizes the advantage of our updated assay over a traditional approach in which nonspecific interactions with the abiotic plate surface likely confounded mucoadhesion measurements. These results offer promise in the delivery and control of drug-release in the emerging field of living microbial therapeutics.

## METHODS

### Microbial strains, plasmids, and growth conditions

All lactobacilli and *Weissella confusa* were cultured in de Man-Rogosa-Sharpe (MRS) medium (RPI Corp, Mt. Prospect, IL) at 37 °C in static microaerobic conditions. *Lactococcus lactis* subsp. cremoris MG1363 was cultured in GM17 medium (M17 broth (BD Difco, Franklin Lakes, NJ), 0.5 % (w/v) glucose) at 30 °C in static microaerobic conditions. *E. coli* strains were cultured in lysogeny broth (LB) (VWR International, Randor, PA) at 37 °C with rotary shaking at 250 rpm. All media was solidified using 1.5 % (w/v) agar (Teknova Inc, Hollister, CA). *E. coli* TG1^*ΔendA*^was used as a host for the construction of the pSIP-based expression vectors listed in Table S2 and cultured as above only supplemented with erythromycin (200 µg/mL) (RPI Corp). When *L. lactis* subsp. cremoris MG1363 was used as hosts for these vectors, the strains were cultured as above only supplemented with erythromycin (5μg/mL).

All cloning was performed in *E. coli* TG1^*ΔendA*^. Beginning with the plasmid pSIP411, the inducible promoter PsppQ^89^ was replaced with the constitutive synthetic promoter P11^90^ using the surrounding sites for restriction endonucleases BglII and NcoI. The genomic DNA of *L. reuteri* ATCC PTA-6475 and *L. reuteri* DSM 17938 were used as templates to amplify the known and putative mucus-associating factors in this study using colony PCR. The pSIP411-P11 backbone, *lr1064*, and *lr0793* were digested with NcoI and XbaI while *cmbA* was digested with NdeI and XbaI. A site mutation was introduced into *nucA* using overlap PCR and the inactive *nucA*^E41Q^ was inserted into a surface-display pSIP411-P11 variant (previously unpublished) by digestion with XhoI and XbaI. This created a fusion construct of secretion signal peptide lp3050^91^, the N-terminal LysM anchoring motif lp3014^91^, and *nucA*^E41Q^. All cloning enzymes were purchased from New England Biolabs (Ipswich, MA).

### Transformation conditions

*L. lactis* MG1363 transformations require GSGM17 medium (GM17, 0.5 M sucrose, 2.5% (w/v) glycine). Electrocompetent cells were prepared by first growing in 5 mL GM17 overnight from a frozen stock optimized for GSGM17 growth. The resulting overnight was subcultured 1:9 in 10 mL GSGM17 and grown overnight. This final overnight culture was subcultured 1:7 in 40 mL GSGM17 and grown 6-7 h (OD_600_ 0.17). Cells were centrifuged at 3000 × g at 4 °C for 25 min and washed once in 40 mL ice-cold EP buffer (0.5 M sucrose, 10 % (w/v) glycerol). The cells were then suspended in 20 mL ice-cold EP buffer and incubated for 15 min on ice. Finally, cells were suspended in 400 µL ice-cold EP buffer and stored at −80 °C in 50 µL aliquots. To transform, 500 ng DNA, always diluted in dH2O to a final volume of 5 µL, was added to 50 µL electrocompetent cells and incubated on ice for 2 min. This mix was transferred to a pre-chilled 2 mm electroporation cuvette and electroporation was performed (2.0 kV, 25 μF, 200 Ω) using a Gene Pulser Xcell (Bio-Rad Laboratories, Hercules, CA) electroporation system. Cells were recovered by immediately adding 950 µL pre-warmed GM17 supplemented with recovery salts (20 mM MgCl_2_, 2 mM CaCl_2_) and incubating statically at 30 °C for 1.5 h. Cells were plated on selective GM17 with erythromycin (5 μg/mL) and grown for at least 24 h at 30 °C.

### Isolation of primary porcine intestinal mucus

The intestines of 8-10-week-old pigs were obtained from a local supplier (Research 87 Inc, Boylston, MA). The small intestine was sectioned into 60 cm segments, squeezed of chyme, and placed in ice-cold phosphate buffered saline (PBS) (137 mM NaCl, 2.7 mM KCl, 10 mM Na_2_HPO_4_, 2 mM KH_2_PO_4_, pH 7.4). Each section was then splayed longitudinally, washed by dipping into a tray of ice-cold PBS, and moved to a second tray of ice-cold PBS. Pinning one end of an intestinal segment, a silicone spatula was used to scrape the mucus layer, placing the mucus in a beaker of ice-cold PBS containing a cOmplete protease inhibitor cocktail tablet (Roche, Basel, Switzerland). The crude PIM was centrifuged at 11000 × g at 4 °C for 10 min, and the supernatant was centrifuged again at 26000 × g at 4 °C for 15 min using an Optima L-90K (Beckman Coulter, Brea, CA) ultracentrifuge with a 70-Ti fixed angle rotor (Beckman Coulter). This clarified PIM supernatant was then lyophilized and stored at 4 °C.

### Plate surface functionalization

The wells of 96-well clear polystyrene U-bottom (Greiner Bio-One, Kremsmünster, Austria) microtiter plates were activated by oxygen plasma etching using a PE-200 (Plasma Etch, Inc. Carson City, NV) industrial benchtop plasma processing system with a R301 RF Generator (Seren IPS, Inc., Vineland, NJ) power supply (105 W, 1 min plasma time, 50 psi vacuum pressure, 20 cc/min O_2_ flow). Immediately, 200 µL of APTES (5% v/v in ethanol) was added to each well, and the plate was sealed with parafilm and incubated at room temperature for 15 min. The plate was then washed twice with ethanol and dried with an air stream, always minimizing exposure to humidity or water. A 1% PIM solution was prepared in PBS. Separately, a 10x solution of EDC (Thermo Fisher Scientific, Waltham, MA) was made in PBS. The EDC was added to the 1% PIM to a final concentration of 5 mM and placed on a rocker for 5 min. During this time, NHS (TCI Chemicals, Portland, OR) was weighed out and added to the 1% PIM to a final concentration of 10 mM, again using a rocker to mix for 5 min. Finally, 200 µL of the PIM/EDC/NHS was added to each well of the APTES functionalized plate. The plate was then sealed with parafilm and incubated at room temperature for 2 h before being left at 4 °C overnight.

### Fluorescent labelling of bacteria

All strains were grown to early stationary phase in 200 mL of their respective growth media (approximately 10 h for LAB strains). To promote flagellum expression *E. coli* was grown for 24 h at 30 °C at 85 rpm. Cells were then centrifuged at 3000 × g at 4 °C for 10 min and washed three times with 50 mL PBS. Cells were then diluted in PBS to OD_600_ 1.0 and labelled by the addition of carboxyfluorescein diacetate (cFDA) (Thermo Fisher Scientific) to a final concentration of 10 μM by incubating at 37 °C for 40 min with rotary shaking at 85 rpm. Again, carboxyfluorescein (cF)-labelled cells were washed three times as above and suspended in 10 mL PBS.

### Mucus adhesion assay

Stained cells were diluted using PBS from the 10 mL stock to OD_600_ values approximately evenly spaced across three orders of magnitude. The mucus solution left after plate functionalization was removed from the microtiter plate by aspiration. Two hundred microliters of each dilution were plated in four technical replicates. A separate 225 µL of each OD_600_ dilution was placed in microfuge tubes, and both the plate and tubes were then incubated at 37 °C for 4 h with orbital shaking at 500 rpm. Using an AquaMax4000 (Molecular Devices, San Jose, CA) plate washer with a cell wash head, the plate was washed three times with PBS (center aspiration 50 µL/well/s; wall dispense 7.2 mL/plate/s; probe height 3 mm) ending with a final aspiration step. The adhered cells were lysed by adding 200 µL lysis buffer (1% (w/v) SDS, 0.1 M NaOH) and incubating at 37 °C for 1 h with rotary shaking at 750 rpm. As a control, each strain was incubated in lysis buffer for 1 h and then plated to check for survival. Even at extremely dense cell concentrations (OD_600_ 10.0), no growth was observed, confirming full cell lysis (data not shown; plates showed no colonies after 48hr). Additionally, the 225 µL cell dilutions were centrifuged at 5000 × g for 5 min and suspended in lysis buffer as fluorescence controls, to relate bulk fluorescence to cell density. All lysates were transferred to a 96-well F-bottom black FluoTrac (Greiner Bio-One) microtiter plate. The plate was passed over with a flame to remove bubbles and fluorescence was measured (Ex/485 nm, Em/520 nm) using a SpectraMax M3 (Molecular Devices) plate reader. Blank wells containing only lysis buffer were used for background subtraction.

The traditional mucus adhesion assay was adopted from MacKenzie, *et al*^38^, Briefly, 200 µL of 0.1% PIM in PBS was added to a 96-well F-bottom black FluoTrac (Greiner Bio-One) microtiter plate and incubated overnight at 4 °C. The plate was then washed three times with PBS, by vacuum aspirating and adding PBS with a multichannel electronic pipette at the slowest dispense speed. The plate was incubated for 1 h at room temperature with Pierce Protein-Free Blocking Buffer (Thermo Fisher Scientific) and again washed with PBS three times. The cF-labelled cells were added at OD_600_ 0.5 and incubated for 4 h at 37 °C under static conditions. After another three washes with PBS, remaining cells were lysed with lysis buffer. Fluorescence was measured as above.

### Adhesion curve fitting and statistical analysis

Plate data was background corrected by subtracting the relative fluorescence units (RFU) of blank wells containing only lysis buffer. The means of experimental replicates were used for further data analysis. By plotting the standards as RFU vs OD_600_, the slope of a linear trend (*C_1_*) was used to associate RFU per unit OD_600_. This relationship, along with the volume-to-surface-area relationship (*V/A*) de-rived from a microtiter plate schematic (Figure S5a) and the CFU/mL per OD_600_ (*C_2_*), can be used to convert values to adhered cells per unit area (*A_y_*):

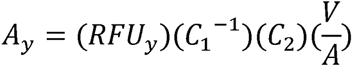

Values for *C_1_* and *C_2_* are summarized in Table S1. A typical sigmoidal (Hill-type) curve was adapted to model fit cell adhesion:

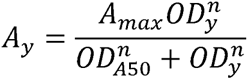

Here, *A_y_* is the adhered cells per unit area at cell concentration *y*, *A_max_* is the maximum possible adhered cells per unit area, OD is the OD_600_ for cell concentration *y*, OD_A50_ is the OD_600_ required for half the maximum number of cells to adhere to the well, and n is the sigmoid factor. The curve parameters were obtained by using the nonlinear curve fitting function in MATLAB (fitnlm) with robust bi-square least squares regression analysis, which applies a weighting factor to discount outliers. Plots of cells per unit area versus OD_600_ were created to visualize the adhesion differences between strains.

## ASSOCIATED CONTENT

The following files are available free of charge.

## SUPPORTING INFORMATION

Additional methods and accompanying figures to confirm mucus adhesion, the expression of mucus-associated proteins, and the specificity of the subsequent mucus binding experiments. A table listing all primers, strains, and plasmids.

- Methods for fluorescence microscopy, ELISA, and Western blot analysis, flow cytometry, and gene knockouts in *E, coli* Nissle 1917
- Figure S1. Adhesion of *L. lactis* MG1363 and *E. coli* Nissle 1917 to different substrates
- Figure S2. Western blots to confirm recombinant expression of mucus-associated factors
- Figure S3. Flow-cytometry confirms surface display of recombinant mucus-associated factors
- Figure S4. Competitive binding of *L. reuteri* and engineered *L. Lactis*
- Figure S5. Mucus characterization after EDC reaction and schematic of Greiner U-bottom plate geometry
- Table S1. Linear relationships between RFU and OD_600_ and OD_600_ and CFU/mL for each strain
- Table S2. List of strains, plasmids, and primers (XLSX) AUTHOR INFORMATION

## AUTHOR INFORMATION

### Author Contributions

Z.J.S.M. performed all experiments. Z.J.S.M. and N.U.N. designed all experiments, analyzed the data, and wrote the manuscript. T.C.C. performed gene deletions in EcN.

## ACKNOWLEDGEMENTS

The authors would like to thank Dr. Huan-Hsuan Hsu and Prof. Xiaocheng Jiang for their advisement and resources when developing the surface immobilization procedure. A special thanks to Josef R. Bober for his advisement with surface-display in LAB.

## FUNDING SOURCES

This work was financially supported by grant numbers R03HD090444 and DP2HD091798 of the National Institutes of Health and a Tufts Collaborates! grant from Tufts University.

## CONFLICTS OF INTEREST

None.

## ABBREVIATIONS

ABC: ATP-binding cassette
APTES: (3-aminopropyl)triethoxysilane
BSA: bovine serum albumin
BscA: cellulose synthase catalytic subunit A
cFDA: carboxyfluorescein diacetate
CmbA: cell and mucus binding protein A
CnBP: collagen binding protein
EcN: *E. coli* Nissle 1917
EDC: *N*-ethyl-*N*’-(3-dimethyl-aminopropyl) carbodiimide
ELISA: enzyme-linked immunosorbent assay
EPS: extracellular polysaccharide
FU: 5-fluorouracil
GI: gastrointestinal
IEC: intestinal epithelial cell
LAB: lactic acid bacteria
MapA: mucus adhesion promoting protein A
MTX: methotrexate
NHS: *N*-hydroxysuccinimide
NucA: nuclease A
PGM: porcine gastric mucus
PIM: porcine intestinal mucus
PTS: proline-threonine/serine
RFU: relative fluorescence unit

## Notes

#### Summary of Updates

Revised Figure 2 New Figures 5, S4, S5

