## Supplemental Tables and Figures for "Quantifying and engineering mucus adhesion of probiotics"

Supplemental Methods

*Fluorescence microscopy, ELISA, and Western blot analysis*

A coupling reaction was prepared as described in the “Plate surface functionalization” section of Methods with a 1% PIM solution combined with EDC/NHS and mixed for 5 min. Sulfo-cyanine5 amine (cya5) (Lumiprobe, Hunt Valley, MA) was added to a final concentration of 1 mg/mL, and the reaction was incubated for 2 h at room temperature, wrapped in foil to prevent light exposure. A Zeba spin desalting column (Thermo Fisher Scientific) was then used to remove excess fluorophore. The mucus-cya5 conjugate was then used in a second coupling reaction, conjugating the mucus to a 96-well clear polystyrene U-bottom (Greiner Bio-One) microtiter plate. After a 2 h incubation, excess volume was aspirated from each well condition. Microscope images were captured using a DMi8 (Leica Microsystems, Inc., Buffalo Grove, IL) microscope with 100x objective. Phase contrast images were taken with 1 ms exposure. Fluorescent images were taken by exciting with a Y5 filter cube with 400 ms exposure and gain set to 4 on LAS X (Leica Microsystems, Inc.) software. Wells were then washed three times with a plate washer as described in the “Mucus adhesion assay” section below.

Microscope images were again taken before an ELISA assay was performed. Each well was blocked for 1 h with cold blocking buffer (5% (w/v) skim milk powder in TBST (20 mM Tris, 150 mM NaCl, 0.1% (v/v) Tween, pH 7.6)). Wells were then washed three times with ice cold TBST, shaking at 150 rpm for 5 min each time. Wells were resuspended in 200 μL blocking buffer containing 0.1 μg rabbit anti-Muc2 (MyBioSource, Inc., San Diego, CA) primary antibody and incubated on ice with shaking at 150 rpm for 1 h. After another three washes, the wells were resuspended in 200 μL blocking buffer containing 0.04 μg goat anti-rabbit IgG-HRP (Abcam, Cambridge, MA) secondary antibody and incubated on ice with shaking at 150 rpm for 30 min. Using a SuperSignal West Femto (Thermo Fischer Scientific) chemiluminescence detection kit, images and luminescence values were recorded after 30 s exposure on a UVP ChemiDoc-It2 810 (Analytik Jena US, LLC., Upland, CA) imager on VisionWorks LS (Analytik Jena US, LLC) analysis software.

*Flow cytometry*

Five milliliter cultures of *L. lactis* MG1363 wild-type or carrying expression vectors were grown overnight as described in the “Microbial strains, plasmids, and growth conditions” section of Methods. Cells were centrifuged at 3000 × g for 5 min and washed twice with PBS before being suspended at OD_600_ 1.0. A 200 μL aliquot was resuspended in 200 μL PBSA (2% (w/v) BSA in PBS) and incubated on ice for 30 min. The cells were then resuspended in 50 μL PBSA containing 1 μg mouse anti-6xHis (Thermo Fisher Scientific) primary antibody and incubated for 1 h at room temperature. The cells were washed three times with PBSA before being suspended in 50 μL PBSA containing 0.4 μg goat anti-mouse IgG-AlexaFluor488 (Thermo Fisher Scientific) secondary antibody and incubated for 15 min on ice, wrapped in foil to prevent light exposure. Finally, the cells were washed three times with PBSA and suspended in 500 μL PBSA. Samples were run on an Attune NxT (Thermo Fisher Scientific) flow cytometer, using a 50 mW blue laser to collect at least 10,000 events. Histograms were analyzed on FCS Express 6 (De Novo Software, Glendale, CA) software.

*Gene knockouts in E. coli Nissle 1917*

Antibiotic cassette insertion and gene knockout (KO) were performed using lambda red recombineering^1^. Kanamycin (kan), chloramphenicol (cat), and ampicillin (amp) were used at 50, 25, and 100 μg/mL, respectively, to maintain plasmids or select for integration of resistance cassettes. Amplification of the cat cassette from pKD3 was used for *bcsA* KO, while amplification of the kan cassette from pKD4 was used for *fliC* KO. Verification of antibiotic resistance cassette integration and removal was verified by Taq colony PCR. Plasmid curing and cassette removal was verified by absence of growth on LB agar plates. All primers for cassette amplification and removal verification are listed in Table S2.

(1)        Datsenko, K. A.; Wanner, B. L. One-Step Inactivation of Chromosomal Genes in Escherichia Coli K-12 Using PCR Products. *PNAS* **2000**, *97* (12), 6640–6645.

Supplemental Figures

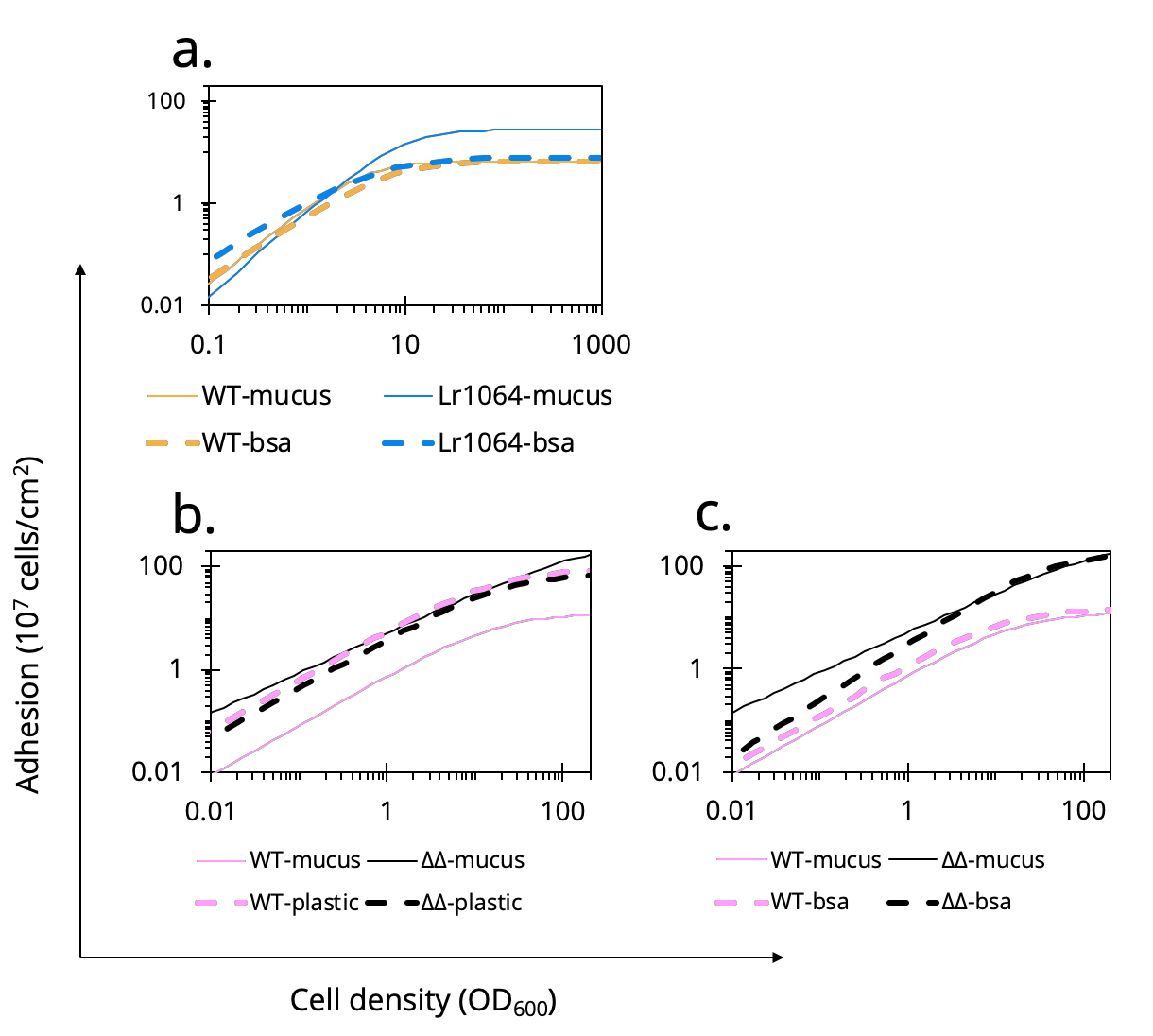

Figure S1. Adhesion of *L. lactis* MG1363 and *E. coli* Nissle 1917 to different substrates

(a.) The observed increase in adhesion of *L. lactis* MG1363 when expressing Lr1064 is shown to be specific to mucus; adhesion does not improve over wild-type *L. lactis* MG1363 when both are adhered to BSA. While the deletion of FliC and BscA increased the ability of EcN to bind to mucus, (b.) the relationship was the reverse when adherence of wild-type EcN and EcN*^ΔfliC,ΔbscA^* was quantified on uncoated polystyrene. (c.) Further investigation shows that EcN*^ΔfliC,ΔbscA^* seems to indiscriminately bind hydrophilic surfaces as adhesion was increased on mucus and BSA. Only the curve fits are shown for clarity.

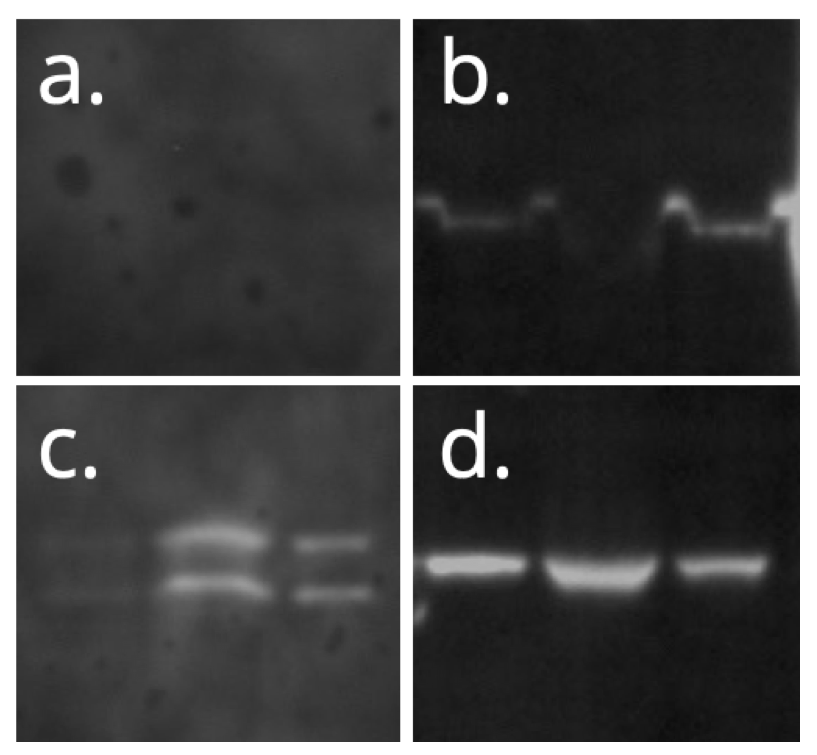

Figure S2. Western blots to confirm recombinant expression of mucus-associated factors

His-tag fused (a.) empty vector, (b.) CmbA, (c.) Lr0793, and (d.) Lr1064 were expressed in *L. lactis* MG1363. Bands show the surface-stripped, soluble, and insoluble fractions. Proteins expressed at the correct size (ladder not shown).

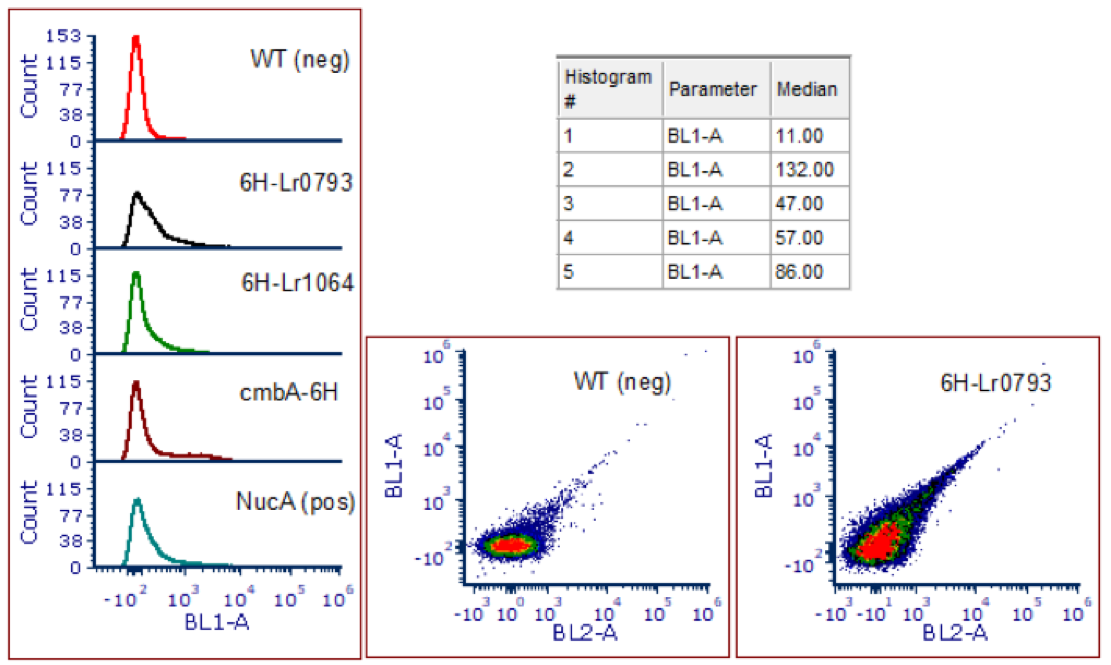

Figure S3. Flow-cytometry confirms surface display of recombinant mucus-associated factors

His-tag fused mucus-associated factors were labelled with AlexaFluor488 conjugated anti-His-tag antibody. Wild-type *L. lactis* MG1363 was used as a negative control and inactivated NucA^E41Q^ was used as a positive control for surface display. A shift to the right on each histogram is an increase in fluorescence, correlated to protein display. Median fluorescence units are shown in the table for each corresponding histogram.

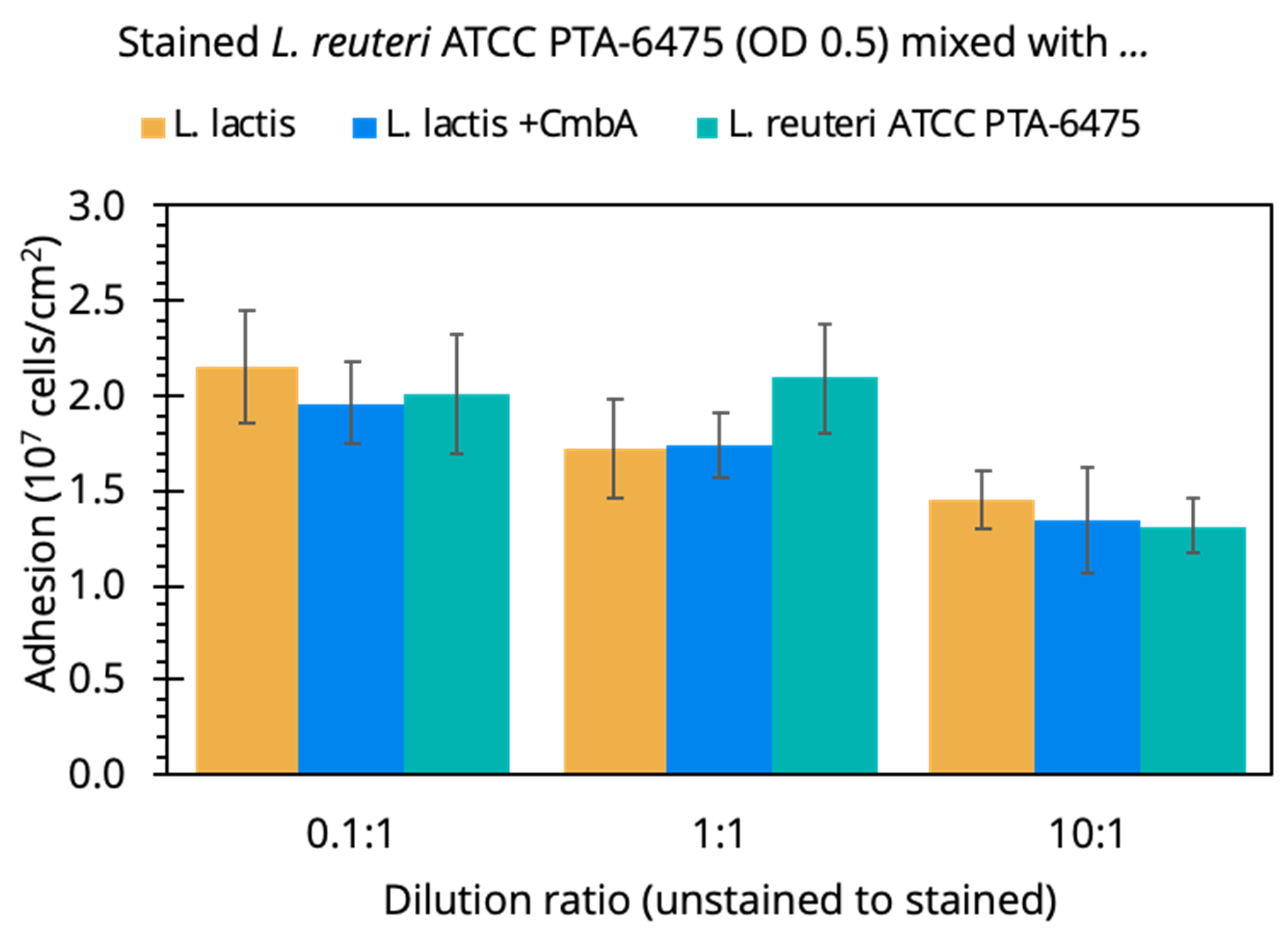

Figure S4. Assessing competitive binding of *L. reuteri* against *L. lactis* (WT & engineered) and itself

*L. reuteri* ATCC PTA-6475 was stained and diluted to OD_600_ 0.5 and mixed at different ratios with unstained cells (WT *L. lactis, L. lactis* + CmbA, or *L. reuteri* ATCC PTA-6475). At low or equal ratios, the binding of stained *L. reuteri* is unchanged regardless of the presence a poor-binding competitor (*L. lactis* MG1363) or a high-binding competitor (*L. lactis* + CmbA or *L. reuteri* ATCC PTA-6475). Only at a 10:1 unstained:stained cell ratio does the binding of *L. reuteri* ATCC PTA-6475 decrease. These cell densities are not representative of the population present in the small intestine, the target of LAB probiotics for therapeutic delivery.

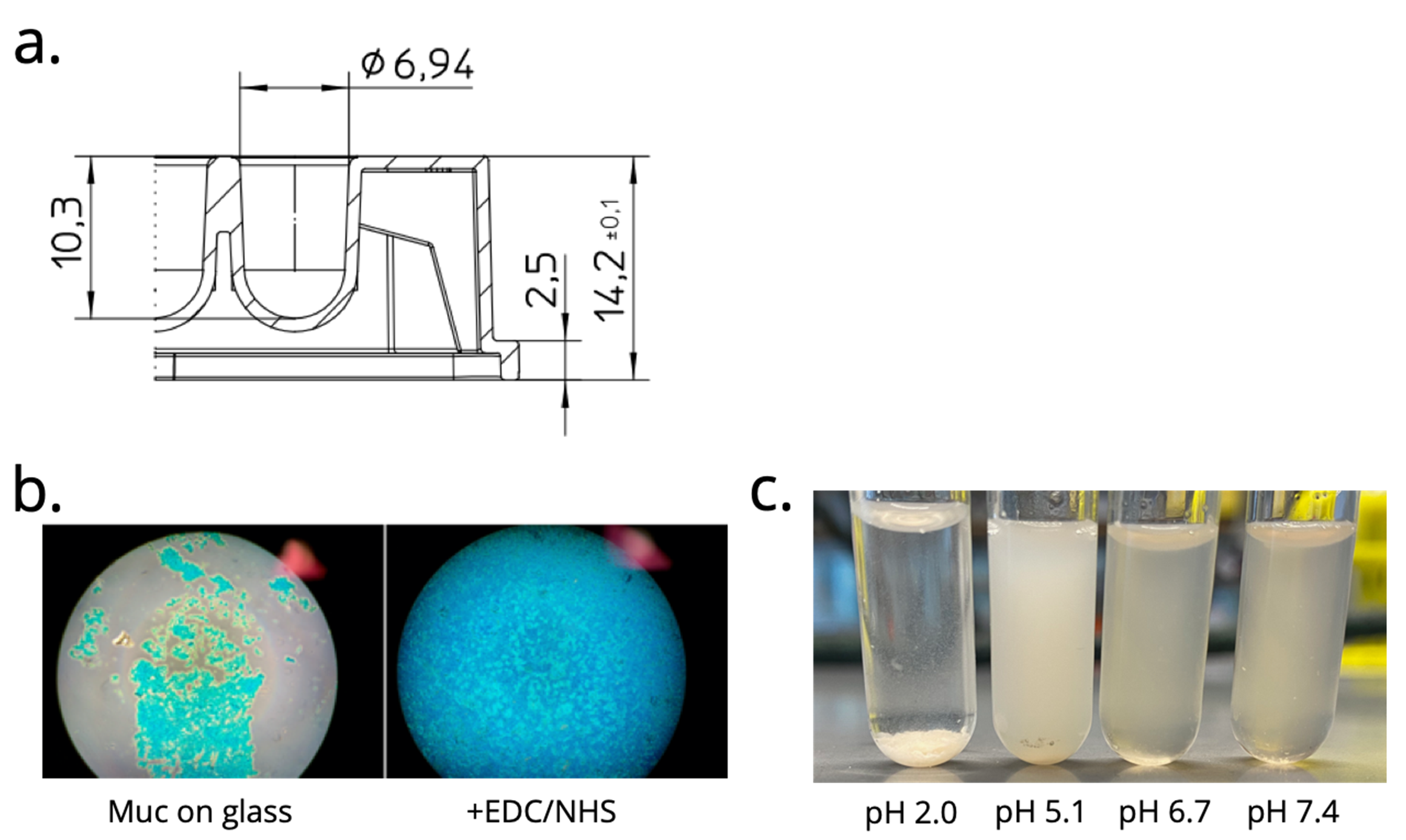

Figure S5. Schematic of Greiner U-bottom plate geometry and mucus characterization after EDC reaction

(a). Sourced from Greiner 96-Well U-bottom product document and used to calculate well surface area. (b.) Visualizing the difference between the traditional (non-specific) and our optimized (covalently-linked) plate functionalization after washing. Mucus was stained with Alcian Blue and viewed using 20x bright-field microscopy. (c.) The EDC modified mucus forms a hydrogel at pH 5.1, consistent with published data, indicating the chemistry does not significantly alter the physical properties of mucus.

Supplemental Tables

Table S1. Linear relationships between RFU and OD_600_ and OD_600_ and CFU/mL for each strain

|  | RFU/OD_600_ | CFU/mL (×10^8^) per OD_600_ |
| --- | --- | --- |
| *L. reuteri* ATCC PTA-5289 | 3109 | 3.7 |
| *L. reuteri* ATCC PTA-6475 | 2794 | 3.3 |
| *L. reuteri* DSM 17938 | 2334 | 6.7 |
| *W. confusa* | 397 | 3.8 |
| *L. rhamnosus* GG | 1002 | 1.4 |
| *E. coli* Nissle 1917 | 51 | 8.0 |
| *L. lactis* MG1363 | 7005 | 6.8 |

Values for CFU/mL acquired by growing cells overnight, checking the OD600 of three dilutions, and plating a 10,000x dilution of those concentrations to count CFUs.

Table S2: see XLSX file.
